# Eyes on the past: Gaze stability differs between temporal expectation and temporal attention

**DOI:** 10.1101/2024.06.07.598015

**Authors:** Aysun Duyar, Marisa Carrasco

**Affiliations:** Department of Psychology, New York University, New York, NY, USA; Center for Neural Science, New York University, New York, NY, USA

**Keywords:** temporal expectation, temporal attention, sequential effects, microsaccades

## Abstract

Temporal expectation and temporal attention distinctly improve performance and gaze stability, and interact at the behavioral and neural levels. Foreperiod—the interval between the preparatory signal and stimulus onset—facilitates temporal expectation. Preceding foreperiod—the foreperiod in the previous trial—modulates expectation at behavioral and oculomotor levels. Here, we investigated whether preceding foreperiod guides temporal attention. Regardless of the preceding foreperiod, temporal attention improved performance, particularly at early moments,and consistently accelerated gaze stability onset and offset by shifting microsaccade timing. However, only with preceding expected foreperiods, attention inhibited microsaccade rates. Moreover, preceding late foreperiods weakened expectation effects on microsaccade rates, but such a weakening was overridden by attention. Altogether, these findings reveal that the oculomotor system’s flexibility does not translate to performance, and suggest that although selection history can be utilized as one of the sources of expectation in subsequent trials, it does not necessarily determine, strengthen, or guide attentional deployment.

## INTRODUCTION

The visual environment is in a continuous state of flux, surpassing the brain’s processing capabilities(Shapiro, 2001). Sequential effects emerge as a consequence of this capacity limitation, such that many perceptual judgements are biased towards previous stimuli in visual (Fischer & Whitney, 2014; Pascucci et al., 2023), auditory (Motala et al., 2020), and olfactory (van der Burg et al., 2022) modalities. Temporal uncertainty –the unpredictability regarding when events may occur– impairs visual processing (Rolke & Hoffman, 2007). The brain’s limited temporal processing capacity emphasizes the necessity for optimizing its resources to navigate the temporal constraints imposed by its own biological architecture (Nobre & van Ede, 2023). *Temporal expectation* –the ability to formulate predictions about when a visual event is likely to occur– and *temporal attention* –the ability to prioritize and selectively process the specific time points based on behavioral relevance regardless of its predictability– are fundamental cognitive mechanisms that act together to distribute these limited resources within a time window (Denison, 2024; Denison, Heeger & Carrasco, 2017; Duyar, Ren & Carrasco, 2024, Duyar, Denison & Carrasco, 2023; Nobre & van Ede, 2023; Todorovic et al., 2015). Together, they improve visual processing based on the probabilities and behavioral goals associated with the dynamic visual environment. It is unknown whether there are sequential effects of event timing on the interplay between temporal attention and expectation.

Temporal expectation improves visual performance, indexed by modulations in accuracy, discriminability, response time, and visual representations (Coull & Nobre, 2008; Cravo et al., 2013; Nobre, Correa & Coull, 2007; Nobre & van Ede, 2023; Rohenkohl et al., 2012, 2014; Shalev & Nobre, 2022; van den Brink et al., 2021; Vangkilde, Coull & Bundesen, 2012). Directing temporal attention to a time point in which the behaviorally relevant stimulus is expected to appear further improves visual performance (Correa, Lupianez, & Tudela, 2005; Denison et al., 2017, 2021; Duyar et al., 2024; Fernández et al., 2019; Griffin, Miniussi, & Nobre, 2001; Jing et al., 2023), at the expense of impairments at unattended but expected moments, resulting in attentional tradeoffs (Denison et al., 2017, 2021; Palmieri & Carrasco, 2024).

Both temporal expectation and attention modulate oculomotor behavior. Our eyes are never completely still, and they move even during fixation. Microsaccades are the fastest and largest of these fixational eye movements, with less than 1 degree of visual angle gaze displacements at a ∼1-2 Hz frequency, and they provide a continuous measure throughout experimental trials (Martinez-Conde et al., 2009; Martinez-Conde, Otero-Millan & Macknik, 2013; Rolfs, 2009; Rucci & Poletti, 2015). Perceptual performance decreases if a microsaccade is executed around the presentation of a brief stimulus (Zuber & Stark, 1966; Beeler, 1967; Hafed & Krauzlis, 2010). Accordingly, microsaccade rate decreases in anticipation of a brief stimulus in visual (Betta & Turatto, 2006; Dankner et al., 2017; Tal-Perry & Yuval-Greenberg, 2020), auditory (Abeles et al., 2020) and tactile (Badde et al., 2020) perception, often leading to improvements in perceptual performance.

Temporal attention further suppresses microsaccade rates in the prestimulus window when temporal attention is deployed to the expected stimulus (Denison, Yuval-Greenberg & Carrasco 2019; Palmieri, Fernández & Carrasco, 2023), although this suppression is not necessarily linked to attentional benefits on visual performance (Palmieri et al., 2023). In addition, the last microsaccade before and the first microsaccade after stimulus onset are executed at an earlier time point with temporal attention as compared to mere expectation (Denison et al., 2019).

Different temporal structures enable distinct types of expectation (Nobre & van Ede, 2018). Foreperiod –the interval between the preparatory signal and the stimulus onset– and preceding foreperiod –the corresponding interval in the previous trial– are two types of temporal structures (Capizzi et al., 2015; Tal-Perry & Yuval-Greenberg, 2023). The foreperiod affects expectations by improving performance and microsaccades: As the foreperiod lengthens and the stimulus is delayed (hazard rate), reaction times decrease, discriminability improves (e.g. Duyar et al., 2024; Janssen & Shadlen, 2005; Los, 2010; Luce, 1991; Schoffelen, Oostenveld, & Fries, 2005), and microsaccade rates diminish (Abeles et al., 2020; Amit et al., 2019; Badde et al., 2020; Dankner et al., 2017). Furthermore, the preceding foreperiod modulates these foreperiod effects on temporal expectation (Drazin, 1961; Los & Van den Heuvel, 2001; Vallesi & Shallice, 2007): As the preceding foreperiod lengthens, the reaction time advantage disappears (Steinborn et al., 2008) and microsaccade inhibition becomes weaker (Tal-Perry & Yuval-Greenberg, 2023).

Research on temporal orienting investigating how cognitive resources are distributed across time, rarely differentiates between temporal expectation and attention when defining and manipulating them. Some studies have disentangled these processes by testing temporal attention at expected timings and showing that temporal attention modulates both performance and oculomotor control beyond temporal expectation (Denison et al., 2017, 2019, 2021; Fernández et al., 2019; Palmieri & Carrasco, 2024; Jing et al., 2023). By keeping temporal expectations constant, these studies have isolated the role of attention. However, in the real world, external noise leads to stochasticity in timing of the events. Strategic allocation of temporal attention relies on the ability to anticipate when behaviorally relevant events are likely to occur (Capizzi et al., 2023; Nobre & van Ede, 2018, 2023). The interplay between attention and expectation under temporal uncertainty has rarely been investigated (Todorovic et al., 2015; Duyar et al., 2023, 2024).

Temporal uncertainty can be quantified and manipulated by systematically increasing the variability of event onsets within a given window (Rolke & Hofmann, 2007). A recent study revealed that this variability influences the allocation of temporal attention. As it increases, attentional benefits on performance decrease at the expected moment—the time point with the highest probability of target occurrence. Moreover, when variability is high, temporal attention tends to be allocated to earlier than the expected moment (Duyar et al., 2024).

The extent of flexibility in attentional deployment under uncertainty, when the target is expected to occur in a temporal window rather than a certain time point, remains unexplored. Here we investigated whether temporal attention is inflexibly allocated to earlier moments of the temporal window, or if recent temporal experiences facilitate trial-to-trial allocation adjustments of temporal attention. Considering the impact of temporal expectations on temporal attention (Duyar et al., 2023, 2024), and of preceding foreperiods on temporal expectations (Capizzi et al., 2015; Los, 2010; Tal-Perry & Yuval-Greenberg, 2022, 2023), we asked whether preceding foreperiods modulate the allocation of temporal attention under uncertainty.

Here we investigated potential sequential effects of the foreperiod on the interaction between attention and expectation on performance and microsaccades. The findings provide insight into how expectations regarding stimulus timing are developed and incorporated into cognitive control of attentional orienting and selection in time.

## METHODS

### Dataset

We reanalyzed behavioral and eye-tracking data collected in a recent study investigating temporal attention modulations on visual performance under temporal uncertainty by Duyar et al. (2024). All observers, apparatus, stimuli, as well as the experimental procedure were identical to those previously reported.

In that study there were 4 levels of temporal precision, manipulated by stimulus timing variability within a session. For this study, we only included the two lowest precision conditions (42% wide and 33% uniform) in the analysis, as in the “hazard rate” section of the previous study (Duyar et al., 2024). To analyze whether temporal attention effects on visual performance and oculomotor dynamics are modulated by preceding foreperiods, we focused on these two temporal precision conditions in which there was high temporal variability and enough number of trials for each type of preceding foreperiod (early_N-1_, expected_N-1_, late_N-1_).

### Observers

The observers were the same sixteen observers (10 females, six males, aged between 22 and 34 years). All participants provided their written consent and had normal or corrected-to-normal vision. The experimental methods complied with the Helsinki Declaration and received approval from the New York University Institutional Review Board.

### Apparatus

Stimuli were generated using an Apple iMac (3.06 GHz, Intel Core 2 Duo) and MATLAB 2012b (Mathworks, Natick, MA, USA), along with the Psychophysics Toolbox (Brainard, 1997; Kleiner et al., 2007), displayed on a color-calibrated CRT monitor (1280×960 resolution, 100 Hz refresh rate). Observers were seated 57 cm away from the screen; their heads were stabilized with a chinrest. Eye position was monitored and recorded using the Eyelink 1000 eye tracker (SR Research, Ottawa, Ontario, Canada). If a participant blinked or moved their eye >1° from the screen’s center during the critical time window (between the precue and the response cue in each trial), the trial was automatically aborted and repeated at the end of the block.

### Stimuli

Stimuli were displayed on a uniform medium gray background. A fixation circle with a diameter of 0.15° of visual angle was presented at the screen center. The placeholders were four small black circles, each 0.2° in size, positioned at the corners of an imaginary square with sides length=2.2°, and remained on the screen throughout the trials to minimize spatial uncertainty.

The target stimuli were 100% contrast, Gaussian-windowed sinusoidal gratings (standard deviation=0.3° 4 cycles per degree (cpd), and random phase). The gratings were oriented either clockwise or counterclockwise from the vertical or horizontal axis. The degree of tilt was independently titrated for each observer and each target interval to attain 75% accuracy at the expected timing in the neutral trials.

The precue and the response cue were auditory stimuli, presented through speakers. The attentional precue was a 200-ms auditory tone, either a sinusoidal wave or a complex waveform composed of sinusoidal waves ranging from 50 to 400 Hz in frequency. A 800 Hz high-frequency sinusoidal tone indicated the first target (T1), a 440 Hz low-frequency tone indicated the second target (T2), and a complex tone was uninformative regarding the target (neutral precue). The same tones, excluding the neutral tone, were also used at the end of the trial as a response cue to indicate observers should respond with regard to the stimulus presented either in T1 or T2.

### Experimental Protocol

Two oriented Gabors were presented sequentially in each trial, and observers performed a two-alternative forced choice (2AFC) orientation discrimination task (**Figure 1A**).

**Figure 1.**
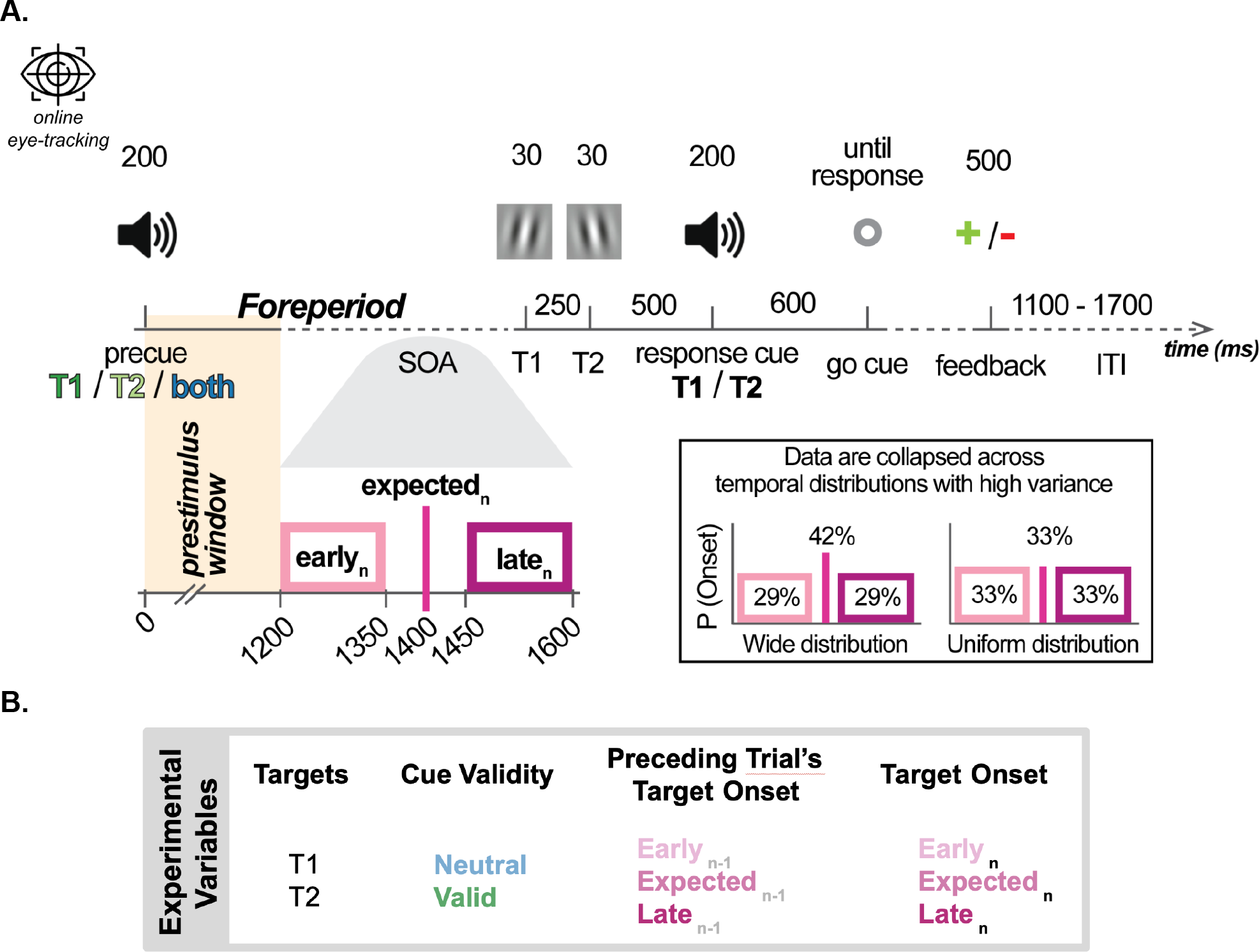
**A**. Psychophysical procedure assessed visual performance at attended and unattended moments, with a variable stimulus timing. The exact timing of the stimulus onset varied across sessions, and we analyzed the data from the sessions with highest variance, “Uniform” and “Wide” temporal distributions. Expected moment is the time point when the stimuli could appear with highest probability (mean point of the temporal distributions). Stimuli could appear earlier or later than the expected time. For microsaccades, we analyzed the “prestimulus window”: the interval between the precue and the earliest possible stimulus onset. Foreperiod is defined as the interval between the precue and T1, and the preceding foreperiod is this interval in the most recent trial. **B**. Summary of experimental variables. There were two targets in each trial, and performance was tested separately for T1 and T2 indicated by the response cue. The precue was either valid or neutral to probe temporal attention. Temporal expectation was manipulated via target timing. Preceding trial’s target onset corresponds to preceding foreperiod, and the target onset corresponds to the current foreperiod.

In the beginning of each trial, an auditory precue instructed participants to attend to either the first target (T1), the second target (T2), or both (neutral). Each target was shown for 30 ms with a stimulus onset asynchrony (SOA) of 250 ms. The response cue was presented 500 ms after the T2 onset, instructing observers to report the orientation of the first (T1), or the second target (T2). Participants responded after the go-cue (800 ms after the response cue onset), signaled by a brightness change of the fixation point.

In neutral trials, the response cue indicated the first or second Gabor patch with equal probability. In valid trials, the pre-cue was 100% accurate, and the response cue consistently matched the pre-cue. Feedback was given after the observer’s response for 500 ms, a red minus for incorrect responses and a green plus for correct responses. Participants were advised to prioritize accuracy rather than speed, and there was no time constraint on their response. The intertrial interval varied from 1100 to 1700 ms.

The onset of the targets with respect to the precue was randomized throughout the experiment. T1 onset varied between 1200-1600 ms and T2 between 1450-1850 ms after the precue, with the most possible onsets occurring at 1400 ms for T1 and 1650 ms for T2 after the precue onset. The stimulus onset asynchrony (SOA) between T1 and T2 was set to 250 ms. This design enabled us to assess performance at the moments both earlier and later than the expected moment.

There were four experimental factors in this study: target, temporal attention, temporal expectation, and preceding foreperiod (**Figure 1B**).

We titrated the neutral performance level at the expected moments (1400 ms for T1 and 1650 ms for T2 following the precue) to a 75% accuracy rate for each target independently, across horizontal and vertical orientations, before starting each session. The initial tilt threshold for each session was established using the Best PEST (Parameter Estimation by Sequential Testing) technique (Lieberman & Pentland, 1982), and this threshold was adjusted after each block as needed to maintain the target neutral accuracy close to 75%.

The trial sequence was randomized for each session, and the sequence of sessions for different temporal precision levels varied across participants. Each participant completed between 5 or 6 sessions, amounting to a total of 5.3 on average per participant (ranging between 2551-3120 trials per participant). Eye tracking data from 9 sessions (out of 85) were corrupted due to unforeseen technical issues with the equipment; thus, we analyzed the eye tracking data from 2252 trials on average per participant (ranging between 1527-3081 trials).

### Behavioral Data Analysis

Data analysis was performed using R software (version 4.2.3; R Core Team, 2023), and repeated measures ANOVA was conducted using the ezANOVA package (version 4.4–0;

Lawrence, 2016) and *η*_*G*_ ^2^ was provided for all F tests. As a rule of thumb, *η*_*G*_ ^2^ = 0.01 is interpreted as a small effect, *η*_*G*_ ^2^ = 0.06 as a medium effect, and *η*_*G*_ ^2^ = 0.14 as a large effect (Cohen, 1988), although this metric is more appropriate to compare across effect sizes in different studies (Lakens, 2013; Thompson, 2007). Greenhouse–Geisser corrections were applied when Mauchly’s test indicated sphericity was violated. To report effect sizes, Cohen’s d was computed for significant differences revealed by post-hoc t-tests, interpreted as d=0.2 as a small effect, d=0.5 as a medium effect, and d=0.8 as a large effect (Cohen, 1988; Fritz, Morris & Richler, 2012).

The discriminability index, d’, was calculated as z(hit rate) - z(false alarm rate) (e.g., Fernández & Carrasco, 2020; Zhang et al., 2019). To prevent infinite values in the calculation of d’, a correction was applied, adding 0.5 to the count of hits, misses, correct rejections, and false alarms (Brown & White, 2005; Hautus, 1995).

Median reaction times (RTs) from the go-cue for correct responses were used in the analysis.

We evaluated behavioral performance at four experimental factors: cue validity (Neutral and Valid), target (T1 and T2), current foreperiod timing (early_N_, expected_N_, late_N_), and preceding foreperiod (early_N-1_, expected_N-1_, late_N-1_).

### Microsaccade preprocessing and analysis

*Preprocessing*. We performed the preprocessing of eye tracking data and microsaccade (MS) analyses using MATLAB. We used a standard velocity-based MS detection algorithm to extract microsaccades (Engbert & Kliegl, 2003). For each eye-tracking sample, eye position displacement was mapped to a 2D velocity-space. Saccades were defined as at least 6 ms duration of consecutive velocities greater than 6 standard deviations from the average velocity within each trial, and microsaccades were defined as saccades smaller than 1 degree of visual angle (dva).

#### Online monitoring and experimental conditions throughout the trial

Microsaccades provide an online measure across the trial Due to the fact that some of the experimental conditions unfold later in the trial, we used different factors in microsaccades analysis than those we used to analyze behavioral performance. Instead of the cue validity and target (Neutral vs Valid for T1 and T2), which are determined by the response cue at the end of the trial, we analyze online attentional modulations based on the precue (T1, T2, Neutral). Similarly, instead of the current foreperiod (early_N_, expected_N_, late_N_), which is unknown to the observer before the stimulus occurs, we collapse across trials with different timings in the prestimulus window. Consequently, we have two factors: precue (T1, T2, Neutral) and preceding foreperiod (early_N-1_, expected_N-1_, late_N-1_) in the prestimulus window. We defined the prestimulus window as the interval from the precue onset to 1200 ms later, which marks the earliest possible stimulus onset (see **Figure 1**).

#### Microsaccade rate analysis

Microsaccade rates were calculated separately for each experimental condition for each observer. We calculated the rate time-course with respect to the precue, and then implemented temporal averaging by using an exponential window causal filter, and lastly smoothing using a moving Gaussian window (σ=100ms). We performed cluster-based permutation tests on the MS rate time courses using MATLAB (Maris & Oostenveld, 2007), and implemented Bonferroni correction.

#### Microsaccade timing analysis

Precise timings of the microsaccades were analyzed to investigate the sequential effects of foreperiod on temporal attention and expectation. We therefore focused on the microsaccades that happen around the stimulus presentation timing. To study prestimulus window MS inhibition (**Figure 1A**), we analyzed the timing of the last MS that occurred before the earliest possible target. We defined inhibition latency as the onset of the last microsaccade occurring in this prestimulus window. For post-stimulus MS rebound, we analyzed the timing of the first MS that occurred after the T1 onset regardless of the trial’s target timing (Bonneh, Adini, & Polat, 2015; Denison, Yuval-Greenberg & Carrasco, 2019).

Trials without any microsaccades were excluded from the timing analysis.

For each observer, we separately detected the inhibitory and rebound MS. We then converted all of the MS onset timings to z-scores, and estimated the kernel density for each precue, expectation, and preceding foreperiod condition. Density was evaluated using 300 equally spaced points between -5 and 5, and the median of each kernel was computed to perform further statistical analyses. We used MATLAB for kernel estimation, and R for statistical tests.

## RESULTS

### Behavior

To investigate sequential effects of stimulus onset on the interaction between temporal expectation and attention, we conducted four-way ANOVAs (2 target x 2 cue validity x 3 current trial’s onset x 3 preceding foreperiod) analyzing discriminability (d’), criterion and reaction time (RT). The experimental factors and the levels were target (T1, T2), attentional cue validity (Valid, Neutral), current foreperiod (Early_N_, Expected_N_, and Late_N_), and preceding foreperiod (Early_N-1_, Expected_N-1_, and Late_N-1_).

For discriminability (**Fig. 2A**) there were significant two-way interactions between target and cue validity (F(1,15) = 12.775, p = 0.003, *η*_*G*_ ^2^*= 0*.*013*), as well as between cue and current trial’s onset (F(2,30) = 3.658, p = 0.038, *η*_*G*_ ^2^ = 0.005). There was a marginal three-way interaction among target, cue validity, and current trial’s onset (F(2,30) = 2.761, p = 0.079, *η*_*G*_ ^2^ *= 0*.*007*). Pairwise t-tests revealed a significant effect of cue validity in the early (p = 0.007, d = 0.313), but not at expected (p > 0.1), or late time points (p > 0.1). There were no main effects or interactions with the preceding foreperiod.

**Figure 2.**
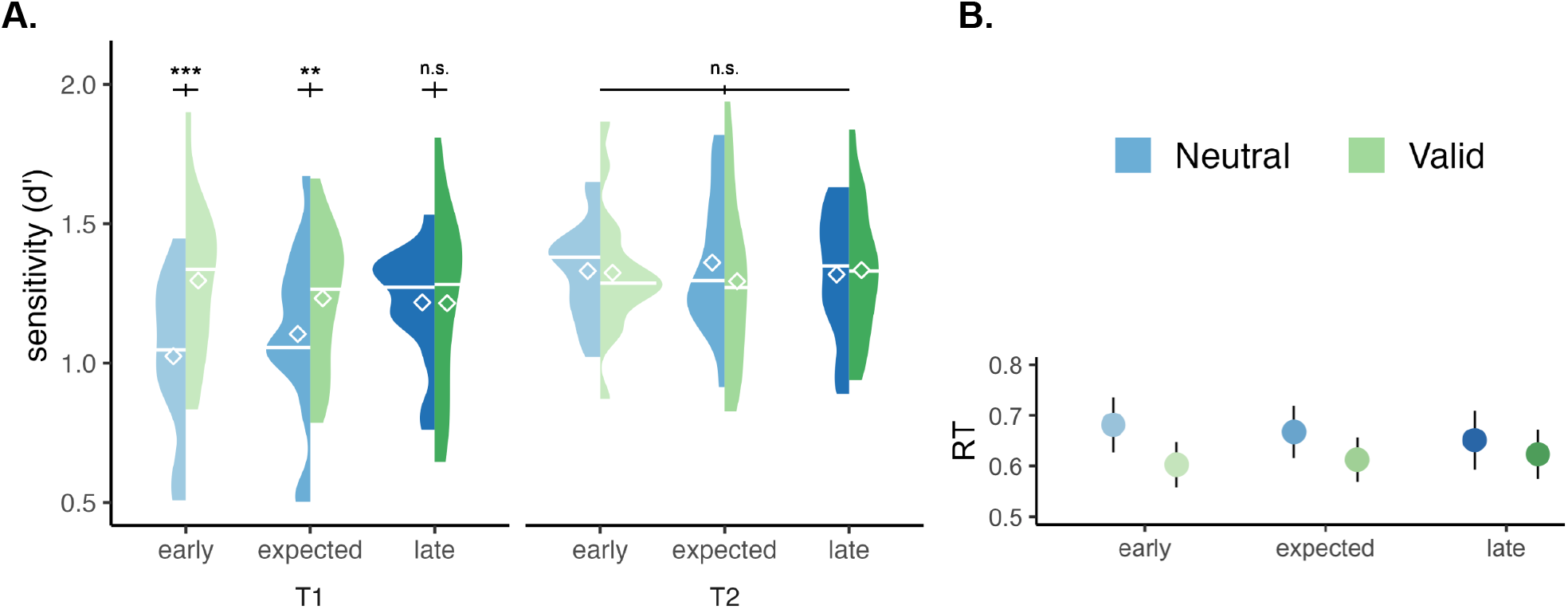
**A**. The effects of target (T1, T2), cue validity (Valid, Neutral), and current trial’s onset timing (Early_N_, Expected_N_, Late_N_) on sensitivity (d’). **B**. The effect of cue validity and current trial’s onset timing on reaction times (RT). Data from Duyar et al., 2024.

For criterion, there were no significant main effects or interactions of any of the experimental factors (all ps > 0.1).

For reaction time (**Fig. 2B**), we found significant main effects of target (F(1,15) = 6.746, p = 0.020, *η*_*G*_ ^2^ = 0.023) and cue validity (F(1,15) = 17.513, p < 0.001, *η*_*G*_ ^2^ = 0.037). There was a two-way significant interaction between target and cue validity (F(1,15) = 17.263, p < 0.001, *η*_*G*_ ^2^ = 0.007), and a marginal three-way interaction among target, cue validity and current trial’s onset (F(2,30) = 0.702, p = 0.0594, *η*_*G*_ ^2^ = 0.001). Pairwise comparisons for the interaction between target and cue validity revealed significant effects of cue validity for both targets, the effect being stronger for T1 (p < 0.001, d = 0.528) than T2 (p < 0.001, d = 0.238).

### Microsaccade rate

We confirmed that peak velocity and amplitude were correlated (R^2^ = 0.913). We analyzed the microsaccade rates in the prestimulus window, defined as the period between the precue onset to 1200 ms, which is the earliest possible stimulus onset (**Figure 1A**).

To investigate attentional effects in this time window, we first compared microsaccade rates within each attentional precue condition, regardless of the preceding foreperiod (**Figure 3A)**. Microsaccade rates in the T1-cued condition were significantly lower than in the neutral condition between 713-1200 ms (487 ms duration, p_Bonferroni_ < 0.01).

**Figure 3.**
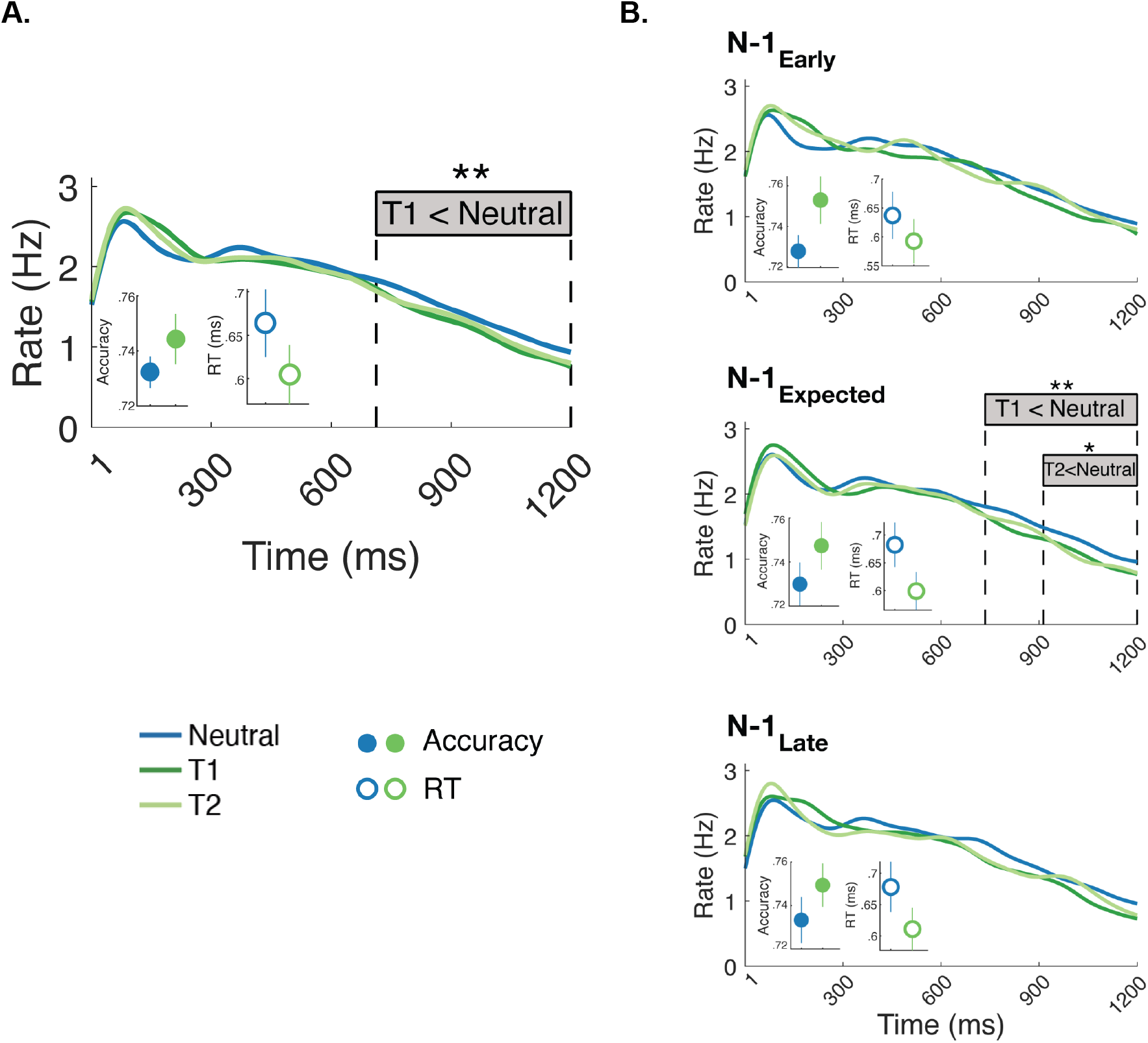
Effects of the attentional precue on microsaccade rates in the prestimulus window. In all graphs, behavioral accuracy increased and RT decreased with temporal attention. Error bars represent standard error of the mean. **A**. Effects of the attentional precue regardless of preceding foreperiod. Microsaccade rates in T1-cued trials were significantly lower than in neutral trials between 713-1200 ms. **B**. Effects of the attentional precue on microsaccade rates within each preceding foreperiod condition. In the Expected_N-1_ condition, neutral trials showed significant differences in microsaccade rates compared to T1-cued trials between 735 and 1200 ms and T2-cued trials between 913 and 1200 ms.

We investigated potential effects of preceding foreperiods on the attentional modulation on microsaccade rates, by comparing the effect of attentional precues (T1, T2, Neutral) on microsaccade rates as a function of the preceding foreperiod (Early_N-1_, Expected_N-1_, and Late_N-1_) (**Figure 3B**). There were two significant clusters in the Expected_N-1_ condition (middle panel), but not in the two other preceding foreperiods (all p_Bonferroni_ > 0.1). Microsaccade rates in the neutral trials were different than the rates in T1-cued trials between 735 and 1200 ms (465 ms duration, p_Bonferroni_ < 0.01), and than in T2-cued trials between 913 and 1200 ms (287 ms duration, p_Bonferroni_ < 0.05).

We then investigated the potential effects of preceding foreperiod in the prestimulus window, regardless of the attentional precue (**Figure 4A**). Microsaccade rates were significantly lower in trials Early_N-1_ than Late_N-1_ between 890 and 1090 ms (200 ms duration, p_Bonferroni_ < 0.05).

**Figure 4.**
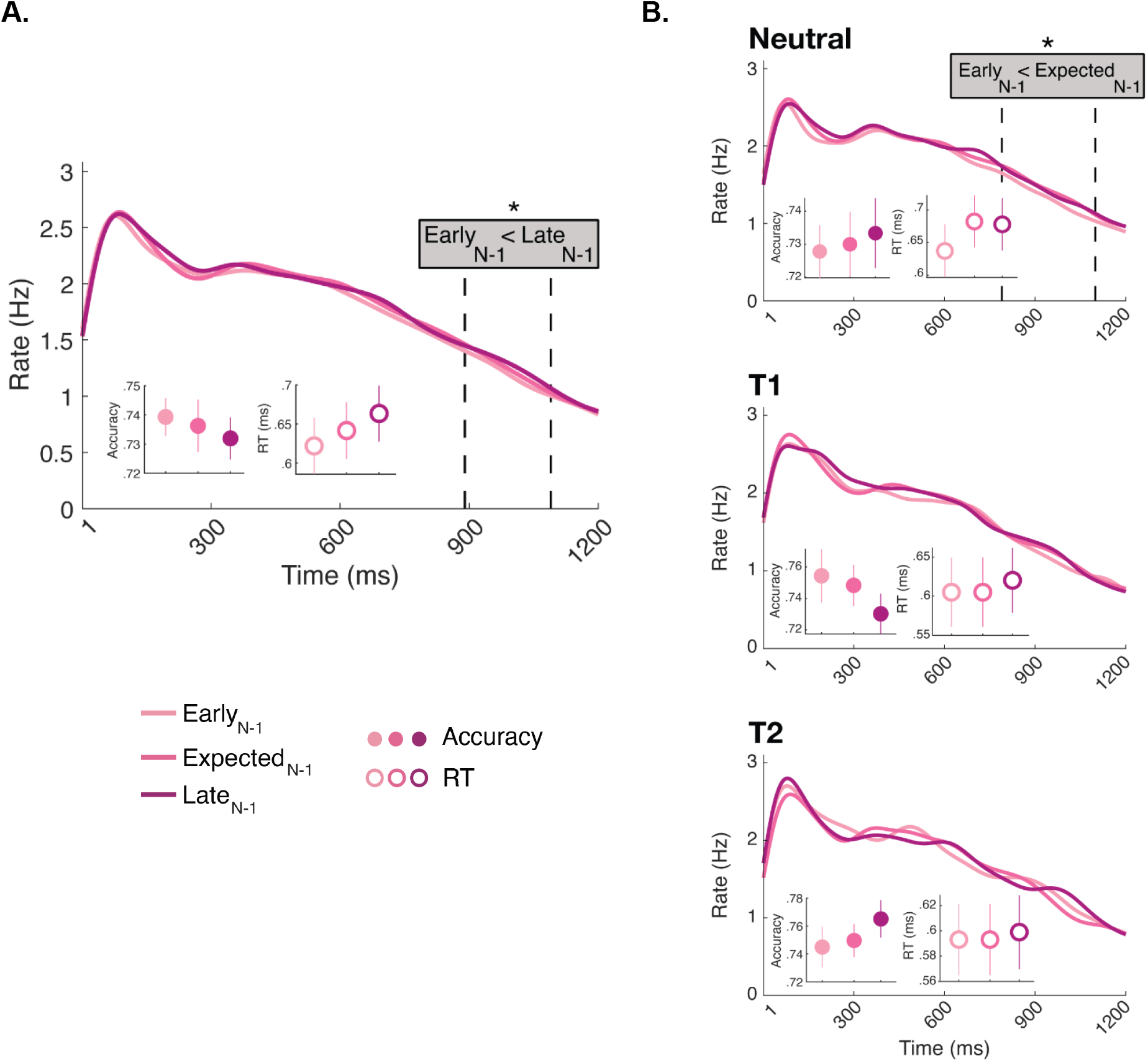
In all graphs, behavioral accuracy and RT plotted as a function of preceding foreperiod (no significant effect). Error bars represent standard error of the mean. **A**. The effect of the preceding foreperiods (Early_N-1_, Expected_N-1_, and Late_N-1_) on microsaccade rates regardless of attentional precue. Early_N-1_ and Late_N-1_ conditions differed between 890 and 1090 ms. **B**.The effect of the preceding foreperiods on microsaccade rates within each attentional precue (Neutral, T1, and T2) condition. In the Neutral condition, a significant difference was observed in the 790-1099 ms interval, highlighting when Early_N-1_ and Expected_N-1_ conditions diverge. No significant effects were found in either T1 or T2 precue conditions.

To investigate whether the effects of preceding foreperiod vary with attention, we compared across microsaccade rates within each attentional precue condition as a function of the preceding foreperiod (Early_N-1_, Expected_N-1_, and Late_N-1_). A significant cluster was detected only in the neutral condition (top panel), where Early_N-1_ and Late_N-1_ microsaccade rates differed between 790-1099 ms after the precue (309 ms duration, p_Bonferroni_ < 0.05; (**Figure 4B**); no significant cluster emerged in the T1 and T2 attentional precue conditions (*p*s > 0.1).

### Microsaccade timing

Two two-way ANOVAs (3 preceding foreperiod x 3 precue) were conducted to analyze their potential timing effects on pre-stimulus inhibition and post-stimulus rebound.

Prestimulus inhibition (**Fig. 5A**) was modulated by precue (F(2, 30) = 9.5958, p = 0.003, *η*_*G*_ ^2^ = 0.187), but not by the preceding foreperiod or its interaction with precue (all ps > 0.1). Pairwise t-tests on precue revealed differences between T1 and T2 (p_holm_ = 0.003, *d* = 0.417), T1 and Neutral (p_holm_ < 0.001, *d* = 1.17)1, and T2 and Neutral (p_holm_ = 0.003, *d* = 0.724). Rebound latency (**Fig. 5B**) was also affected by precue (F(2, 30) = 3.8838, p = 0.0495, *η*_*G*_ ^2^ = 0.0470), but not by the preceding foreperiod or its interaction with precue (all ps>0.1). Pairwise t-tests on the precue revealed a significant difference between T1 and Neutral (p_holm_ = 0.018, *d* = 0.533), and a marginal effect between T2 and Neutral (p_holm_ = 0.072, *d* = 0.316).

**Figure 5.**
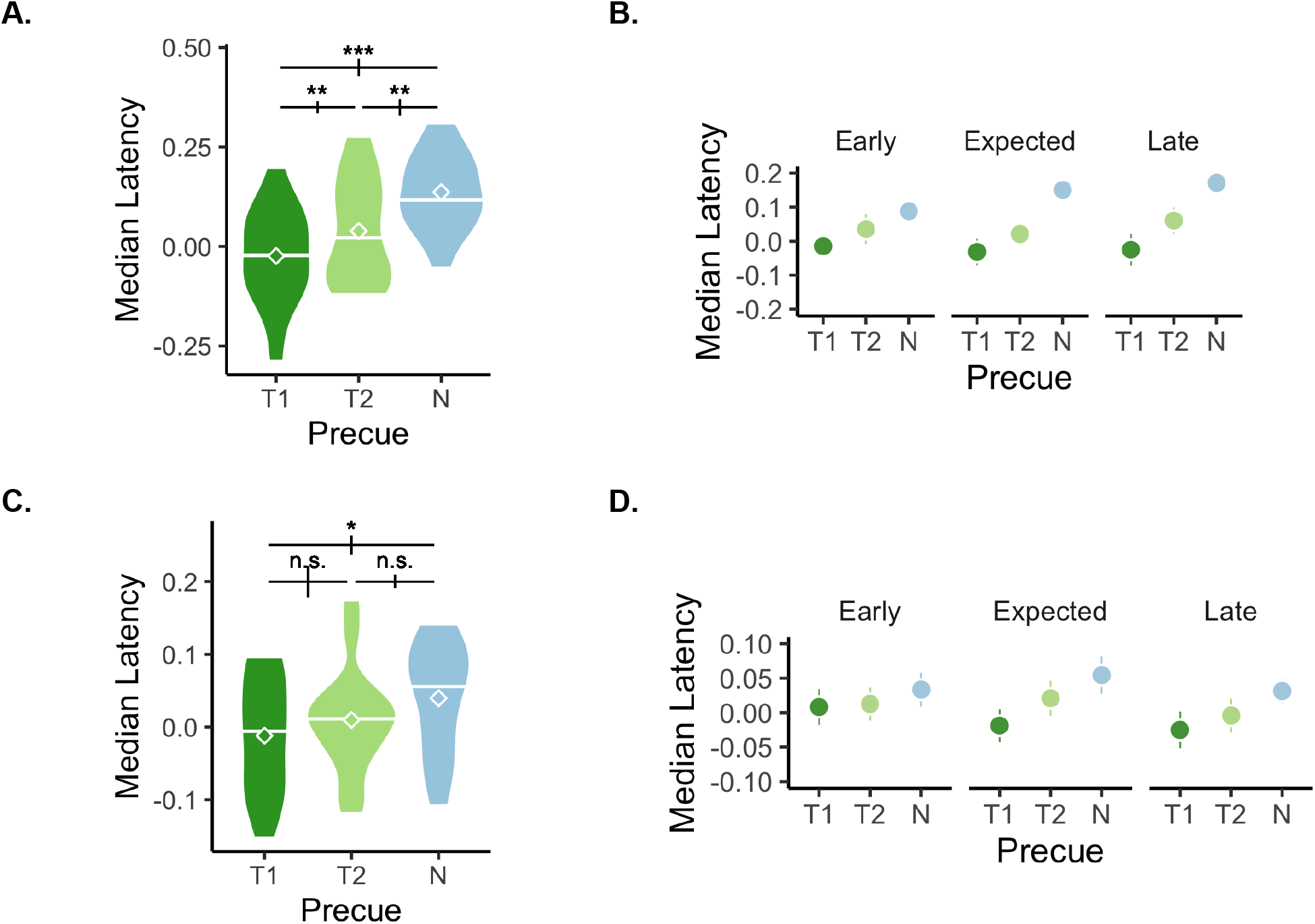
Microsaccades timing around stimulus onset. **A**. Inhibition of microsaccades was analyzed within the prestimulus window (from precue onset to 1200 ms). There was a main effect of precue, but no effect of preceding foreperiod. **B**. Inhibition latency shown separately for preceding foreperiod. The effect of precue on microsaccade latency was consistent across preceding foreperiod. **C**. Microsaccade rebound after the stimulus onset revealed a main effect of precue, and no effect of preceding foreperiod. **D**. The effect of precue on rebound latency was consistent across preceding foreperiod.

## DISCUSSION

Temporal expectation, the ability to predict the onset of an upcoming event, and attention, the ability to select task-relevant specific time points, work together to distribute limited resources. In this study, we investigated the flexibility of temporal attention under temporal uncertainty on behavioral performance and oculomotor dynamics. We reanalyzed data from Duyar et al.’s (2024) study showing that temporal uncertainty results in high attentional benefits on performance at earlier than expected moments, which gradually decrease with target delay. We assessed the potential role of preceding foreperiods on attentional deployment in time. We found that the preceding foreperiod does not modulate the effects of temporal attention on either behavioral performance or microsaccade timing, but it does modulate microsaccade rates. Therefore, although the perceptual system represents the preceding foreperiod in oculomotor dynamics, it does not affect the specific time when temporal attention benefits performance.

The effects of temporal attention on performance were stable (**Fig. 3**). This finding is noteworthy given that many perceptual judgements are biased towards previous stimuli in visual (Cicchini et al., 2014, 2023; Fischer & Whitney, 2014; Pascucci et al., 2023; Taubert et al., 2016), auditory (Motala et al., 2020) and olfactory (van der Burg et al., 2022) modalities. The preceding foreperiod affects the speed of perceptual judgements (Capizzi et al., 2015; Possamai et al., 1973; Steinborn & Langner, 2012; Tal-Perry & Yuval-Greenberg, 2022). Reaction times decrease when the current foreperiod is shorter than the preceding foreperiod, indicating that temporal expectation can be flexibly updated on a trial-by-trial basis. Given the established effects of trial sequence in perception and response time, we hypothesized that the preceding foreperiod could affect allocation of temporal attention under temporal uncertainty. However, we found that the benefits of temporal attention under uncertainty remained consistent regardless of preceding foreperiod (**Fig. 3**). This finding indicates that control of temporal attention is not updated based on the timing of individual trials, and suggests that observers aim to optimize their overall visual performance.

Temporal attention modulated the precise timing of microsaccades around the target presentation: It shifted both the last microsaccade before the stimulus onset and the first microsaccade after the stimulus offset earlier, the effects being stronger for T1 than T2 under temporal uncertainty (**Fig. 5**). These findings agree with those in a similar study, but in which T1 and T2 always occurred with predictable timings (Denison et al., 2019). In addition to being present under temporal uncertainty, this earlier suppression and rebound were also independent of the preceding foreperiod. The earlier suppression in the prestimulus window, helping gaze stabilization, seems resilient to varying temporal dynamics and immediate past history.

We also investigated whether the preceding foreperiod modulates temporal attention effects on microsaccade rate in the prestimulus window. Temporal attention strengthens the prestimulus oculomotor inhibition beyond mere expectation (Denison et al., 2019; Palmieri et al., 2023). Both of these studies investigated temporal attention with no temporal uncertainty; they kept the stimulus timing constant to control for temporal expectation. Here we found that this is the case even when there is temporal uncertainty (**Fig. 3A**) and such strengthening depends on the preceding foreperiod: Temporal attention facilitates prestimulus rate inhibition only in trials that follow a trial with expected timing (**Fig. 3B**).

With regard to expectation, prestimulus inhibition was stronger for early than preceding late foreperiods across attention conditions (**Fig. 4A**). This was also the case for the trials in the neutral condition (**Fig. 4B**, top panel), consistent with a recent finding on temporal expectation (Tal-Perry & Yuval-Greenberg, 2023). Interestingly, when observers were instructed to selectively attend to T1 or T2 such a preceding foreperiod effect did not emerge (**Fig. 4B**, middle and bottom panels). These findings indicate that attention overrode the preceding foreperiod effect that occurs with expectation.

Repetition of the same experimental condition can improve performance (e.g., Asgeirsson, Kristjánsson, & Bundesen, 2014; Becker, 2008; Chun & Nakayama, 2000). Researchers propose prior experience, or “selection history” as a distinct type of attentional control, interpreting this performance improvement as an involuntary attraction of attention based on recent experience (Anderson, 2024; Awh, Belopolsky & Theeuwes, 2012). However, studies investigating sequential effects in various types of attention have yielded mixed results regarding selection history effects on attentional deployment (e.g. Choe & Kim, 2022; Kristjánsson & Ásgeirsson, 2019; Ramgir & Lamy, 2021; White, Rolfs & Carrasco, 2013; Yashar et al., 2017). For instance, it modulates the deployment of presaccadic attention at the saccade target (White, Rolfs & Carrasco, 2013), whereas in visual search, selection history does not influence stimulus-driven spatial attention (Yashar et al., 2017).

Our current behavioral findings –consistent attentional deployment regardless of temporal selection history– suggest that although selection history can be utilized as one of the sources of expectation in subsequent trials, it does not necessarily determine, strengthen, or guide attentional deployment.

In conclusion, oculomotor control and visual performance can be dissociated under temporal uncertainty. Temporal attention allocation is independent from the preceding foreperiod for both behavioral performance and precise timing of microsaccades, but not for microsaccade rates. Preceding foreperiod effects for microsaccade rates in expectation (neutral trials) were overridden by temporal attention. Altogether, these findings suggest that microsaccade rates are not a reliable index of temporal attention allocation and its benefits on visual performance, and while preceding temporal experiences are taken into consideration by the oculomotor system, the influence of the most recent foreperiod is not robust enough to drive attentional allocation under uncertainty.

## ACKNOWLEDGEMENTS

We thank Rania Ezzo, Simran Purokayastha, Helena Palmieri and Mrugank Dake for helpful comments. This research was funded by the National Eye Institute (R01-EY019693 to M.C.).

## REFERENCES

Abeles, D., Amit, R., Tal-Perry, N., Carrasco, M., & Yuval-Greenberg, S. (2020). Oculomotor inhibition precedes temporally expected auditory targets. Nature Communications, 11(1). 10.1038/s41467-020-17158-9

Amit, R., Abeles, D., Carrasco, M., & Yuval-Greenberg, S. (2019). Oculomotor inhibition reflects temporal expectations. NeuroImage, 184, 279–292, doi:10.1016/j.neuroimage.2018.09.026

Anderson, B. A. (2024). Trichotomy revisited: A monolithic theory of attentional control. Vision Research, 217, 108366. 10.1016/j.visres.2024.108366

Ásgeirsson, Á. G., Kristjánsson, Á., & Bundesen, C. (2014). Independent priming of location and color in identification of briefly presented letters. Attention, Perception, & Psychophysics, 76(1), 40–48. 10.3758/s13414-013-0546-6

Awh, E., Belopolsky, A. V., & Theeuwes, J. (2012). Top-down versus bottom-up attentional control: A failed theoretical dichotomy. Trends in Cognitive Sciences, 16(8), 437–443. 10.1016/j.tics.2012.06.010

Badde, S., Myers, C. F., Yuval-Greenberg, S., & Carrasco, M. (2020). Oculomotor freezing reflects tactile temporal expectation and aids tactile perception. Nature Communications, 11(1): 3341, doi:10.1038/s41467-020-17160-1

Becker, S. I. (2008). Can intertrial effects of features and dimensions be explained by a single theory? Journal of Experimental Psychology: Human Perception and Performance, 34(6), 1417–1440. 10.1037/a0011386

Beeler, G. W. (1967). Visual threshold changes resulting from spontaneous saccadic eye movements. Vision Research, 7(9–10), 769–775. 10.1016/0042-6989(67)90039-9

Betta, E., & Turatto, M. (2006). Are you ready? I can tell by looking at your microsaccades. NeuroReport, 17(10), 1001–1004. 10.1097/01.wnr.0000223392.82198.6d

Brainard D. H. (1997) The psychophysics toolbox. Spatial Vision, 10, 433–436

Bonneh, Y. S., Adini, Y., & Polat, U. (2015). Contrast sensitivity revealed by microsaccades. Journal of Vision, 15(9), 11. 10.1167/15.9.11

Brown, G. S., & White, K. G. (2005). The optimal correction for estimating extreme discriminability. Behavior Research Methods, 37(3), 436–449. 10.3758/bf03192712

Capizzi, M., Correa, Á., Wojtowicz, A., & Rafal, R. D. (2015). Foreperiod priming in temporal preparation: Testing current models of sequential effects. Cognition, 134, 39–49, 10.1016/j.cognition.2014.09.002

Capizzi, M., Martín-Signes, M., Coull, J. T., Chica, A. B. & Charras, P. (2023). A transcranial magnetic stimulation study on the role of the left intraparietal sulcus in temporal orienting of attention. Neuropsychologia 184, 108561. 10.1016/j.neuropsychologia.2023.108561

Chun, M. M., & Nakayama, K. (2000). On the functional role of implicit visual memory for the adaptive deployment of attention across scenes. Visual Cognition, 7(1-3), 65–81. 10.1080/135062800394685

Choe, E., & Kim, M.-S. (2022). Eye-specific attentional bias driven by selection history. Psychonomic Bulletin & Review, 29(6), 2155–2166. 10.3758/s13423-022-02121-0

Cicchini, G. M., Anobile, G., & Burr, D. C. (2014). Compressive mapping of number to space reflects dynamic encoding mechanisms, not static logarithmic transform. Proceedings of the National Academy of Sciences, 111(21), 7867–7872. 10.1073/pnas.1402785111

Cicchini, G. M., Mikellidou, K., & Burr, D. C. (2023). Serial Dependence in Perception. Annual Review of Psychology, 75(1), 129–154. 10.1146/annurev-psych-021523-104939

Cohen, J. (1988). Statistical power analysis for the behavioral sciences (2nd ed.). Hillsdale, NJ: Erlbaum.

Correa, Á., Lupiáñez, J., & Tudela, P. (2005). Attentional preparation based on temporal expectancy modulates processing at the perceptual level. Psychonomic Bulletin & Review, 12(2), 328–334. 10.3758/BF03196380

Coull, J., & Nobre, A. (2008). Dissociating explicit timing from temporal expectation with fmri. Current Opinion in Neurobiology, 18(2), 137–144. 10.1016/j.conb.2008.07.011

Cravo, A. M., Rohenkohl, G., Wyart, V., & Nobre, A. C. (2013). Temporal expectation enhances contrast sensitivity by phase entrainment of low-frequency oscillations in visual cortex. Journal of Neuroscience, 33(9), 4002–4010. 10.1523/jneurosci.4675-12.2013

Dankner, Y., Shalev, L., Carrasco, M., & Yuval-Greenberg, S. (2017). Prestimulus inhibition of saccades in adults with and without attention-deficit/hyperactivity disorder as an index of temporal expectations. Psychological Science, 28(7), 835–850. 10.1177/0956797617694863

Denison, R. N. (2024). Visual temporal attention from perception to computation. Nature Reviews Psychology, 3(4), 261–274. 10.1038/s44159-024-00294-0

Denison, R. N., Carrasco, M., & Heeger, D. J. (2021). A dynamic normalization model of temporal attention. Nature Human Behaviour, 5(12), 1674–1685. 10.1038/s41562-021-01129-1

Denison, R. N., Heeger, D. J., & Carrasco, M. (2017). Attention flexibly trades off across points in time. Psychonomic Bulletin & Review, 24(4), 1142–1151. 10.3758/s13423-016-1216-1

Denison, R. N., Yuval-Greenberg, S., & Carrasco, M. (2019). Directing voluntary temporal attention increases fixational stability. Journal of Neuroscience, 9(39), 353–363,

Drazin, D. H. (1961). Effects of foreperiod, foreperiod variability, and probability of stimulus occurrence on simple reaction time. Journal of Experimental Psychology, 62(1), 43–50. 10.1037/h0046860

Duyar, A., Denison, R. N., & Carrasco, M. (2023). Exogenous temporal attention varies with temporal uncertainty. Journal of Vision, 23(3), 9. 10.1167/jov.23.3.9

Duyar, A., Ren, S., & Carrasco, M. (2024). When temporal attention interacts with expectation. Scientific Reports, 14(1). 10.1038/s41598-024-55399-6

Engbert, R., & Kliegl, R. (2003). Microsaccades uncover the orientation of covert attention. Vision Research, 43(9), 1035–1045. 10.1016/s0042-6989(03)00084-1

Fernández, A., & Carrasco, M. (2020). Extinguishing exogenous attention via transcranial magnetic stimulation. Current Biology, 30(20). 10.1016/j.cub.2020.07.068

Fernández, A., Denison, R. N., & Carrasco, M. (2019). Temporal attention improves perception similarly at foveal and parafoveal locations. Journal of Vision, 19(1), 12. 10.1167/19.1.12

Fischer, J., & Whitney, D. (2014). Serial dependence in visual perception. Nature Neuroscience, 17(5), 738–743. 10.1038/nn.3689

Fritz, C. O., Morris, P. E., & Richler, J. J. (2012). Effect size estimates: Current use, calculations, and interpretation. Journal of Experimental Psychology: General, 141(1), 2–18. 10.1037/a0024338

Griffin, I. C., Miniussi, C., & Nobre, K. (2001). Orienting attention in time. SSRN Electronic Journal,6(1), 660–671. 10.2139/ssrn.4048588

Hafed, Z. M., & Krauzlis, R. J. (2010). Microsaccadic suppression of visual bursts in the primate superior colliculus. Journal of Neuroscience, 30(28), 9542–9547. 10.1523/jneurosci.1137-10.2010

Hautus, M. J. (1995). Corrections for extreme proportions and their biasing effects on estimated values OFD′. Behavior Research Methods, Instruments, & Computers, 27(1), 46–51. 10.3758/bf03203619

Janssen, P., & Shadlen, M. N. (2005). A representation of the hazard rate of elapsed time in Macaque Area LIP. Nature Neuroscience, 8(2), 234–241. 10.1038/nn1386

Jing, C., Jin, H., Li, W., Wu, Z., Chen, Y., & Huang, D. (2023). Temporal attention affects contrast response function by response gain. Frontiers in Human Neuroscience, 16. 10.3389/fnhum.2022.1020260

Kleiner, M., Brainard, D.H., & Pelli, D.G. (2007). What’s new in Psychtoolbox-3? Perception, 36(14), 1–16.

Kristjánsson, Á., & Ásgeirsson, Á. G. (2019). Attentional priming: Recent insights and current controversies. Current Opinion in Psychology, 29, 71–75. 10.1016/j.copsyc.2018.11.013

Lakens, D. (2013). Calculating and reporting effect sizes to facilitate cumulative science: A practical primer for T-tests and ANOVAs. Frontiers in Psychology, 4. 10.3389/fpsyg.2013.00863

Lieberman H. R., & Pentland A.P. (1982) Microcomputer-based estimation of psychophysical thresholds: The Best PEST. Behavior Research Methods & Instrumentation, 14, 21–25.10.3758/BF03202110

Los, S. A. (2010). Foreperiod and sequential effects: Theory and Data. Attention and Time, 289–302. 10.1093/acprof:oso/9780199563456.003.0021

Los, S. A., & Van Den Heuvel, C. E. (2001). Intentional and unintentional contributions of nonspecific preparation during reaction time foreperiods. Journal of Experimental Psychology: Human Perception and Performance, 27(2), 370–386. 10.1037//0096-1523.27.2.370

Luce, R. D. (1986). Response times: Their role in Inferring Elementary Mental Organization,Oxford Psychology Series (New York, 1991; online edn, Oxford Academic, 1 Jan. 2008), 10.1093/acprof:oso/9780195070019.001.0001

Maris, E., & Oostenveld, R. (2007). Nonparametric statistical testing of EEG- and Meg-Data. Journal of Neuroscience Methods, 164(1), 177–190. 10.1016/j.jneumeth.2007.03.024

Martinez-Conde, S., Macknik, S. L., Troncoso, X. G., & Hubel, D. H. (2009). Microsaccades: a neurophysiological analysis. Trends in Neurosciences, 32(9), 463–475. 10.1016/j.tins.2009.05.006

Martinez-Conde, S., Otero-Millan, J., & Macknik, S. L. (2013). The impact of microsaccades on Vision: Towards a unified theory of saccadic function. Nature Reviews Neuroscience, 14(2), 83–96. 10.1038/nrn3405

Motala, A., Zhang, H., & Alais, D. (2020). Auditory rate perception displays a positive serial dependence. I-Perception, 11(6), 204166952098231. 10.1177/2041669520982311

Nobre, A., Correa, A., & Coull, J. (2007). The hazards of Time. Current Opinion in Neurobiology, 17(4), 465–470. 10.1016/j.conb.2007.07.006

Nobre, A. C., & van Ede, F. (2018). Anticipated moments: Temporal structure in attention. Nature Reviews Neuroscience, 19(1), 34–48, doi:10.1038/nrn.2017.141

Nobre, A. C., & van Ede, F. (2023). Attention in Flux. Neuron, 111(7), 971–986. 10.1016/j.neuron.2023.02.032

Palmieri, H., & Carrasco, M. (2024). Task demand mediates the interaction of spatial and temporal attention. Scientific Reports, 14(1). 10.1038/s41598-024-58209-1

Palmieri, H., Fernández, A. & Carrasco, M. (2023). Microsaccades and temporal attention at different locations of the visual field. Journal of Vision, 23(5), 6. 10.1167/jov.23.5.6

Pascucci, D., Tanrikulu, Ö. D., Ozkirli, A., Houborg, C., Ceylan, G., Zerr, P., Rafiei, M., & Kristjánsson, Á. (2023). Serial dependence in visual perception: A Review. Journal of Vision, 23(1), 9. 10.1167/jov.23.1.9

Possamai, C. A., Granjon, M., Requin, J., & Reynard, G. (1973). Sequential effects related to foreperiod duration in simple reaction time. Perceptual and Motor Skills, 36(3, Pt. 2), 1185–1186. 10.2466/pms.1973.36.3c.1185

Ramgir, A., & Lamy, D. (2021). Does feature intertrial priming guide attention? The jury is still out. Psychonomic Bulletin & Review, 29(2), 369–393. 10.3758/s13423-021-01997-8

Rohenkohl, G., Cravo, A. M., Wyart, V., & Nobre, A. C. (2012). Temporal expectation improves the quality of sensory information. Journal of Neuroscience, 32(24), 8424–8428. 10.1523/jneurosci.0804-12.2012

Rohenkohl, G., Gould, I. C., Pessoa, J., & Nobre, A. C. (2014). Combining spatial and temporal expectations to improve visual perception. Journal of Vision, 14(4), 8–8. 10.1167/14.4.8

Rolfs M. (2009). Microsaccades: small steps on a long way. Vision Research, 49(20), 2415–2441. 10.1016/j.visres.2009.08.010

Rolke, B., & Hofmann, P. (2007). Temporal uncertainty degrades perceptual processing. Psychonomic Bulletin & Review, 14(3), 522–526. 10.3758/bf03194101

Rucci, M., & Poletti, M. (2015). Control and Functions of Fixational Eye Movements. Annual Review of Vision Science, 1, 499–518. 10.1146/annurev-vision-082114-035742

Schoffelen, J.-M., Oostenveld, R., & Fries, P. (2005). Neuronal coherence as a mechanism of effective corticospinal interaction. Science, 308(5718), 111–113. 10.1126/science.1107027

Shalev, N., & Nobre, A. C. (2022). Eyes wide open: Regulation of arousal by temporal expectations. Cognition, 224, 105062. 10.1016/j.cognition.2022.105062

Shapiro, K. (Ed.). (2001). The limits of attention: Temporal constraints in human information processing. Oxford University Press. 10.1093/acprof:oso/9780198505150.001.0001

Steinborn, M. B., & Langner, R. (2012). Arousal modulates temporal preparation under increased time uncertainty: Evidence from higher-order sequential foreperiod effects. Acta Psychologica, 139(1), 65–76, 10.1016/j.actpsy.2011

Steinborn, M. B., Rolke, B., Bratzke, D., & Ulrich, R. (2008). Sequential effects within a short foreperiod context: Evidence for the conditioning account of temporal preparation. Acta Psychologica, 129(2), 297–307. 10.1016/j.actpsy.2008.08.005

Tal-Perry, N., & Yuval-Greenberg, S. (2022). The spatiotemporal link of temporal expectations: Contextual temporal expectation is independent of spatial attention. Journal of Neuroscience, 42(12), 2516–2523.

Tal-Perry, N., & Yuval-Greenberg, S. (2023). Sequential effect and temporal orienting in prestimulus oculomotor inhibition. Journal of Vision, 23(14), 1. 10.1167/jov.23.14.1

Taubert, J., Van der Burg, E., & Alais, D. (2016). Love at second sight: Sequential dependence of facial attractiveness in an on-line dating paradigm. Scientific Reports, 6(1). 10.1038/srep22740

Thompson, B. (2007). Effect sizes, confidence intervals, and confidence intervals for effect sizes. Psychology in the Schools. 44, 423–432.

Todorovic, A., Schoffelen, J.-M., van Ede, F., Maris, E., & de Lange, F. P. (2015). Temporal expectation and attention jointly modulate auditory oscillatory activity in the beta band. PLOS ONE, 10(3). 10.1371/journal.pone.0120288

Vallesi, A., & Shallice, T. (2007). Developmental dissociations of preparation over time: Deconstructing the variable foreperiod phenomena. Journal of Experimental Psychology: Human Perception and Performance, 33(6), 1377–1388. 10.1037/0096-1523.33.6.1377

van den Brink, R. L., Murphy, P. R., Desender, K., de Ru, N., & Nieuwenhuis, S. (2020). Temporal expectation hastens decision onset but does not affect evidence quality. The Journal of Neuroscience, 41(1), 130–143. 10.1523/jneurosci.1103-20.2020

Van der Burg, E., Toet, A., Brouwer, A.-M., & van Erp, J. B. (2022). Sequential effects in odor perception. Chemosensory Perception, 15(1), 19–25. 10.1007/s12078-021-09290-7

Vangkilde, S., Coull, J. T., & Bundesen, C. (2012). Great expectations: Temporal expectation modulates perceptual processing speed. Journal of Experimental Psychology: Human Perception and Performance, 38(5), 1183–1191. 10.1037/a0026343

White, A. L., Rolfs, M., & Carrasco, M. (2013). Adaptive deployment of spatial and feature-based attention before saccades. Vision Research, 85, 26–35. 10.1016/j.visres.2012.10.017

Yashar, A., White, A. L., Fang, W., & Carrasco, M. (2017). Feature singletons attract spatial attention independently of feature priming. Journal of Vision, 17(9), 7. 10.1167/17.9.7

Zhang, H., Morrone, M. C., & Alais, D. (2019). Behavioural oscillations in visual orientation discrimination reveal distinct modulation rates for both sensitivity and response bias. Scientific Reports, 9(1). 10.1038/s41598-018-37918-4

Zuber, B. L., & Stark, L. (1966). Saccadic suppression: Elevation of visual threshold associated with saccadic eye movements. Experimental Neurology, 16(1), 65–79. 10.1016/0014-4886(66)90087-2

